# Alpha-1-antitrypsin binds to the glucocorticoid receptor with biological significance in macrophages

**DOI:** 10.1101/2022.03.11.483896

**Authors:** Xiyuan Bai, An Bai, Michele Tomasicchio, James R. Hagman, Ashley M. Buckle, Arnav Gupta, Vineela Kadiyala, Shaun Bevers, Karina A. Serban, Kevin Kim, Zhihong Feng, Kathrin Spendier, Guy Hagen, Lorelenn Fornis, David E. Griffith, Monika Dzieciatkowska, Robert A. Sandhaus, Anthony N. Gerber, Edward D. Chan

## Abstract

Alpha-1-antitrypsin (AAT), a serine protease inhibitor produced mainly by the liver, is the third most abundant protein in plasma. While a canonical receptor for AAT has not been identified, AAT can be internalized into the cytoplasm and is known to affect gene regulation. Since AAT has significant anti-inflammatory properties affecting many cell types including macrophages, we examined whether AAT binds the cytoplasmic glucocorticoid receptor (GR) in macrophages. We report the novel finding that AAT binds to GR in macrophages using several approaches, including co-immunoprecipitation, mass spectrometry, microscale thermophoresis, and molecular modeling. The mass spectrometry data are available *via* ProteomeXchange with identifier PXD030989. We further demonstrate that AAT induction of angiopoietin-like 4 protein and AAT inhibition of lipopolysaccharide-induced nuclear factor-kappa B activation and interleukin-8 production are mediated, in part, through AAT–GR interaction. Furthermore, this interaction contributes to a host-protective role against mycobacteria in macrophages. The interaction of AAT and GR described in this study identifies a mechanism for the antiinflammatory and host-defense properties of AAT.

## INTRODUCTION

Severe alpha-1-antitrypsin (AAT) deficiency most commonly occurs in individuals who possess the protease inhibitor (Pi) ZZ genotype, resulting in a substantially increased risk of precocious emphysema. Other disorders associated with PiZZ include bronchiectasis, panniculitis, cirrhosis, and anti-neutrophil cytoplasmic antibody-mediated vasculitis (de Serres & Blanco, 2014; Parr *et al*, 2007; Strnad *et al*, 2020). Additionally, AAT has potent antiinflammatory properties and is host-protective against HIV, influenza, *Pseudomonas aeruginosa*, and non-tuberculous mycobacteria (Bai *et al*, 2019; Bryan *et al*, 2010; Chan *et al*, 2007; Harbig *et al*, 2020; Pott *et al*, 2013; Shapiro *et al*, 2001; Whitney *et al*, 2011; Zhou *et al*, 2016; Zhou *et al*, 2011); however, the molecular basis of these AAT-associated activities is not well characterized.

AAT resides in the cytoplasm of monocytes and macrophages and may be produced by them (Barbey-Morel *et al*, 1987). Alternatively, elastase-bound AAT may enter cells from the extracellular milieu through endocytosis by low-density lipoprotein receptor-related protein-1 (LRP1) or through clathrin-coated vesicles and caveolae-mediated endocytosis (Serban & Petrache, 2016; Sohrab *et al*, 2009; Zhou *et al*, 2015). The scavenger receptor B type I has also been proposed as a mechanism for the endocytosis of AAT (Lockett *et al*, 2015). In the cytoplasm, an anti-inflammatory effect of AAT occurs *via* binding to IkBa (nuclear factor of kappa light polypeptide gene enhancer in B-cells inhibitor, alpha), the basal inhibitor of nuclear factor-kappa B (NFκB). Binding of AAT to IkBa prevents IkBa phosphorylation, ubiquitination, and targeted proteasome degradation (Zhou *et al.*, 2011). Through these actions, AAT inhibits NFκB activation. Additional proposed anti-inflammatory functions of AAT include binding of interleukin-8 (IL-8) (Bergin *et al*, 2010) and inhibiting the ability of ADAM-17 (<u>A</u> <u>D</u>isintegrin <u>A</u>nd <u>M</u>etalloprotease <u>17</u>) to cleave membrane-bound tumor necrosis factor (TNF) to soluble TNF (de Queiroz *et al*, 2020). Via its function as a serine protease inhibitor (serpin), AAT also irreversibly binds and inhibits thrombin, plasmin, and caspase-3, with the lattermost interaction preventing the apoptosis of endothelial cells (Gans & Tan, 1967; Petrache *et al*, 2006; Serban & Petrache, 2016).

Independent of its serine protease inhibitory activity, the mechanism(s) underlying anti-inflammatory and disease-mitigating properties of AAT remains poorly understood. It is well-established that AAT regulates transcription and is found in the nucleus (Aggarwal *et al*, 2016; Janciauskiene *et al*, 2018); however, its ability to translocate into the nucleus is unknown. In contrast, the pleiotropic actions of the glucocorticoid (GC)-glucocorticoid receptor (GR) complex have been extensively studied (Necela & Cidlowski, 2004). GR regulates pro- and anti-inflammatory genes by binding to transcriptional activators, co-activators, repressors, and co-repressors (Necela & Cidlowski, 2004). In the absence of ligand, GR resides in the cytoplasm bound to the chaperones Hsp90 and Hsp70. Upon binding to its canonical ligand, cortisol, the chaperone molecules dissociate from GR, and the activated receptor enters the nucleus to inhibit or induce transcription. GR signaling in response to GC leads to both anti-inflammatory effects and immune suppression. A known connection between AAT and GR is that GR activation by GC increases AAT production at the transcriptional level (Kadiyala *et al*, 2016; Ni *et al*, 2017).

Given that both AAT and GCs have anti-inflammatory properties and reside in the cytoplasmic compartment, we investigated the possibility that AAT interacts directly with GR. We found that AAT binds GR and that this molecular interaction has anti-inflammatory and anti-mycobacterial functions in macrophages. Furthermore, this AAT–GR interaction lends itself to manipulation within the context of the anti-inflammatory, immunosuppressive, and immune-enhancing activities of the GR.

## RESULTS

### AAT co-localizes with GR

Fluorescent immunocytochemical staining of macrophages was performed using anti-GR and anti-AAT antibodies conjugated to different fluorochromes to determine whether AAT and GR are expressed under basal conditions. Both AAT and the GR co-localized in the cytoplasm and, to a lesser extent, within the nuclei of differentiated THP-1 macrophages by confocal microscopy (**Figure 1A**). MDM from an individual who possesses two normal M alleles for AAT (PiMM) and RAW 264.7 macrophages also showed that AAT and GR are found intracellularly by fluorescent microscopy (**Figures 1B** and **1C**, respectively). Similarly, AAT and GR expression in human macrophages were confirmed by immunoblotting whole-cell lysates (**Figure 1D**).

**Figure 1.**
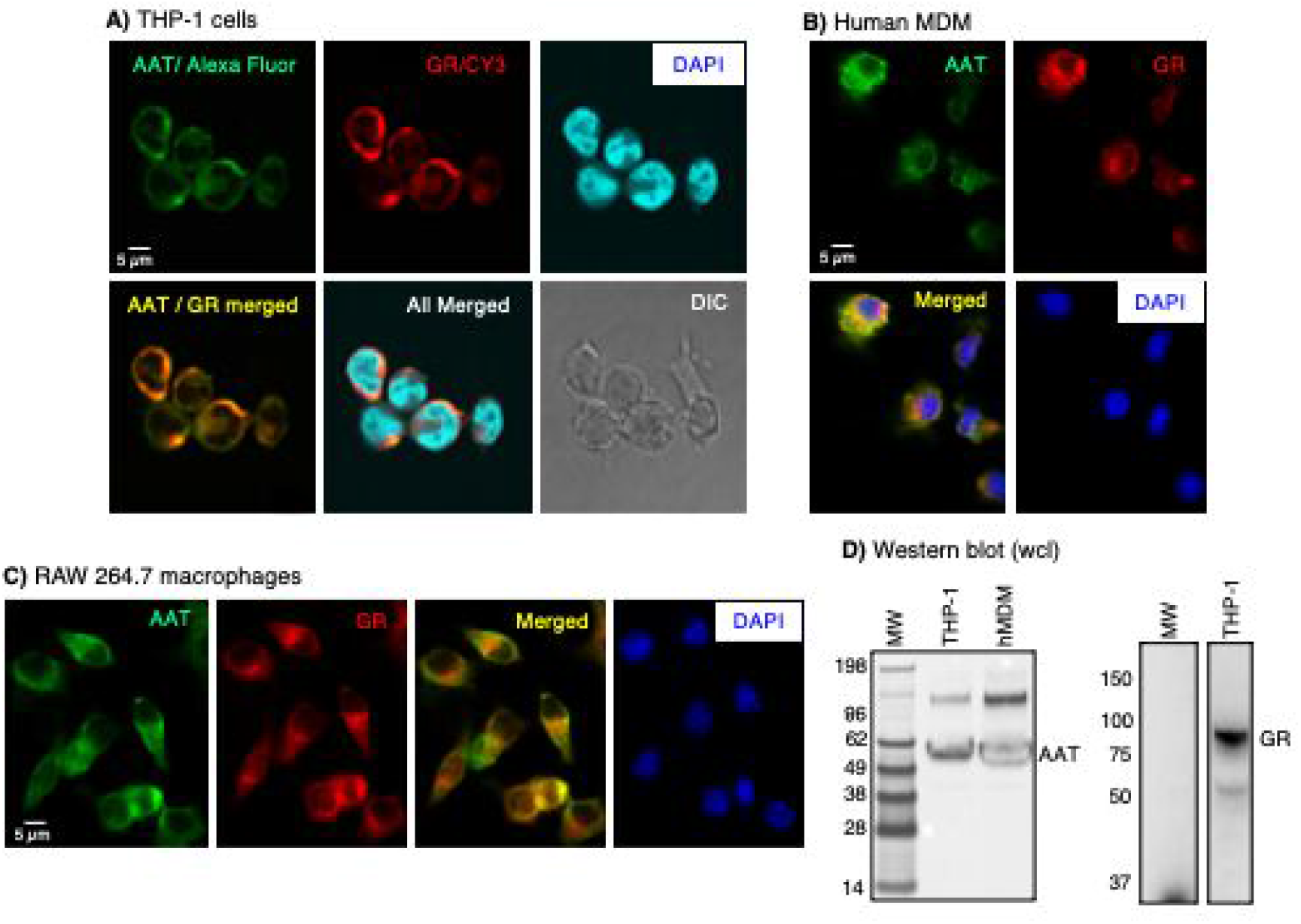
Immunofluorescence staining for glucocorticoid receptor (GR) and alpha-1-antitrypsin (AAT). **(A)** Human THP-1 differentiated macrophages, **(B)** human monocyte-derived macrophages (MDM), and **(C)** RAW 264.7 murine macrophages were immunostained using anti-GR and anti-AAT antibodies conjugated to the fluorochromes Cy3 and Alexa fluor, respectively. THP-1 macrophages were imaged with a confocal microscope. Human monocyte-derived macrophages (MDM) and murine RAW 264.7 macrophages were imaged with a fluorescent microscope. The scale bars represent 5 μm. **(D)** Immunoblotting for AAT and GR in human macrophages. Images shown are representative of three independent experiments.

### AAT binds to the GR

#### Co-immunoprecipitation-immunoblot approach

To investigate whether AAT binds to the GR in differentiated THP-1 macrophages, wholecell lysates were immunoprecipitated (IP’d) for GR with polyclonal anti-GR or non-immune IgG control antibodies using protein A sepharose beads. The beads were washed, centrifuged, the bound proteins extracted, separated by SDS-PAGE, and immunoblotted with an anti-AAT polyclonal antibody. In contrast to non-immune IgG, immunoprecipitation (IP) for the GR followed by immunoblotting for AAT revealed a distinct band at ~52 kDa, which is consistent with the molecular mass of pharmacologic AAT (Glassia^®^) and AAT from whole-cell lysates (**Figure 2A**). Inversely, cell lysates were IP’d for AAT and the separated proteins were immunoblotted for GR. As shown in **Figure 2B**, GR was detected in the fraction IP’d for AAT but not in the fraction IP’d with non-immune IgG.

**Figure 2.**
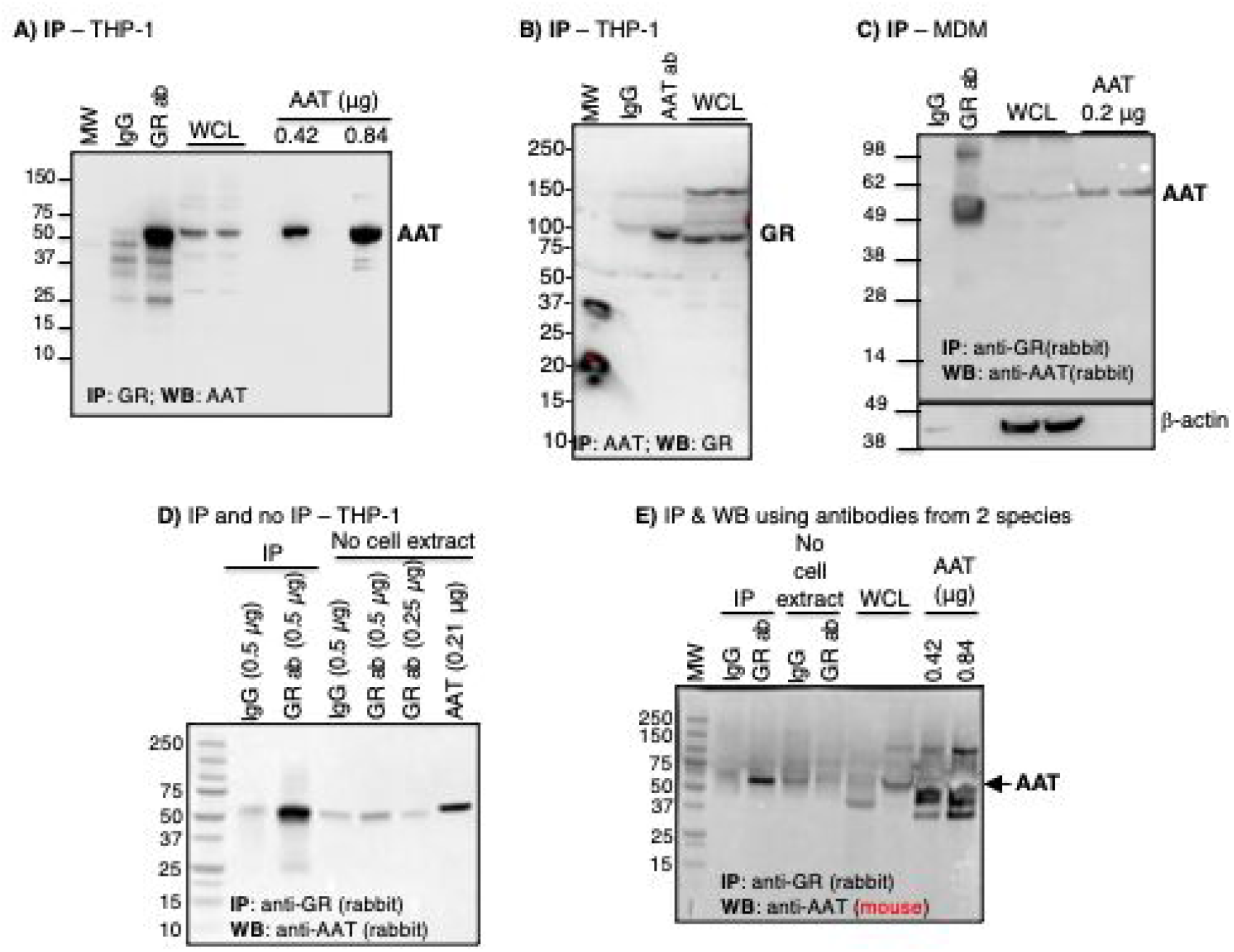
Alpha-1-antitrypsin (AAT) co-immunoprecipitates with glucocorticoid receptor (GR). **(A)** THP-1 macrophage lysates were immunoprecipitated (IP’d) with anti-GR polyclonal antibody, followed by immunoblotting of the IP’d fraction to detect AAT. **(B)** THP-1 macrophage lysates were IP’d with anti-AAT polyclonal antibody, followed by immunoblotting of the IP’d fraction for GR. **(C)** Human monocyte-derived macrophages (MDM) were IP’d with anti-GR polyclonal antibody, followed by immunoblotting of the IP’d fraction for AAT. **(D)** Comparing the relative amount of IgG and anti-GR antibodies (alone, “No cell extract”) used in the immunoprecipitation (IP) experiments, with THP-1 macrophage lysates IP’d with non-immune IgG or anti-GR, followed by immunoblotting for AAT. **(E)** THP-1 macrophage lysates were IP’d with anti-GR polyclonal antibodies followed by immunoblotting of the IP’d fraction with an anti-AAT monoclonal antibody. **ab**=antibody; **WB**=western blot; **WCL**=whole cell lysates. Lanes labeled “No cell extract” contains only IgG or anti-GR antibody alone.

To corroborate the interaction between AAT and GR in primary human macrophages, MDMs were prepared from an individual with the PiMM AAT phenotype. Compared to the non-immune IgG control, IP of MDM lysates with anti-GR revealed the presence of AAT within the coprecipitates (**Figure 2C**). In summary, the data provide evidence that AAT and the GR form a complex in THP-1 macrophages and MDM.

We further confirmed the integrity of the AAT–GR binding interaction by excluding an experimental artifact resulting from the molecular weight similarity between heavy chain immunoglobulins and AAT. We employed two approaches. In the first approach, the THP-1 macrophage lysates were IP’d as previously indicated with the anti-GR polyclonal antibody. In addition, 0.5 μg of anti-GR or non-immune IgG antibodies – the amount used for IP in each condition – were loaded onto separate wells of the SDS-PAGE gel. While the migration of IgG heavy chain was similar to that of AAT, the intensities of the relevant bands in the lanes that contained only either anti-GR or non-immune IgG (with no IP of cell extract) were significantly lower than the IP’d band, indicating that the prominent band at 52 kDa following IP for GR mostly represents AAT and not the heavy chain of IgG (**Figure 2D**). In the second approach, we conducted IP with rabbit anti-GR polyclonal antibody as before but used a mouse anti-AAT monoclonal antibody for immunoblotting. This approach excluded detection of the anti-GR antibody (or the non-immune IgG) used for the IP in the immunoblot. This strategy still demonstrated a prominent band at 52 kDa (representing co-IP’d AAT) following IP with anti-GR and immunoblotting for AAT but not with the control IgG (**Figure 2E**).

#### Co-IP mass spectrometry approach

To further validate the physical interaction between AAT and the GR, we adopted a mass spectrometry approach. THP-1 macrophage lysates were incubated with anti-GR or non-immune IgG as before, and the IP’d proteins were separated with SDS-PAGE under reducing conditions. After electrophoresis, the gel was stained with Coomassie blue. Protein bands that corresponded in molecular weight to AAT were excised from the gel and prepared for mass spectrometry (**Figure 3A, dashed and solid-line black boxes**). Since the molecular weight of AAT may vary based on its glycosylation, we also excised an adjacent area of the gel just beneath the first excised site to ensure that an adequate amount of purported AAT was retrieved (**Figure 3A, dashed and solid line red boxes**). Mass spectrometry of the band that corresponded to pharmacologic control AAT (Glassia^®^) (**Figure 3A, dotted black box, AAT lane**) revealed 1762 total peptides and 62 unique peptides that referenced to AAT.

**Figure 3.**
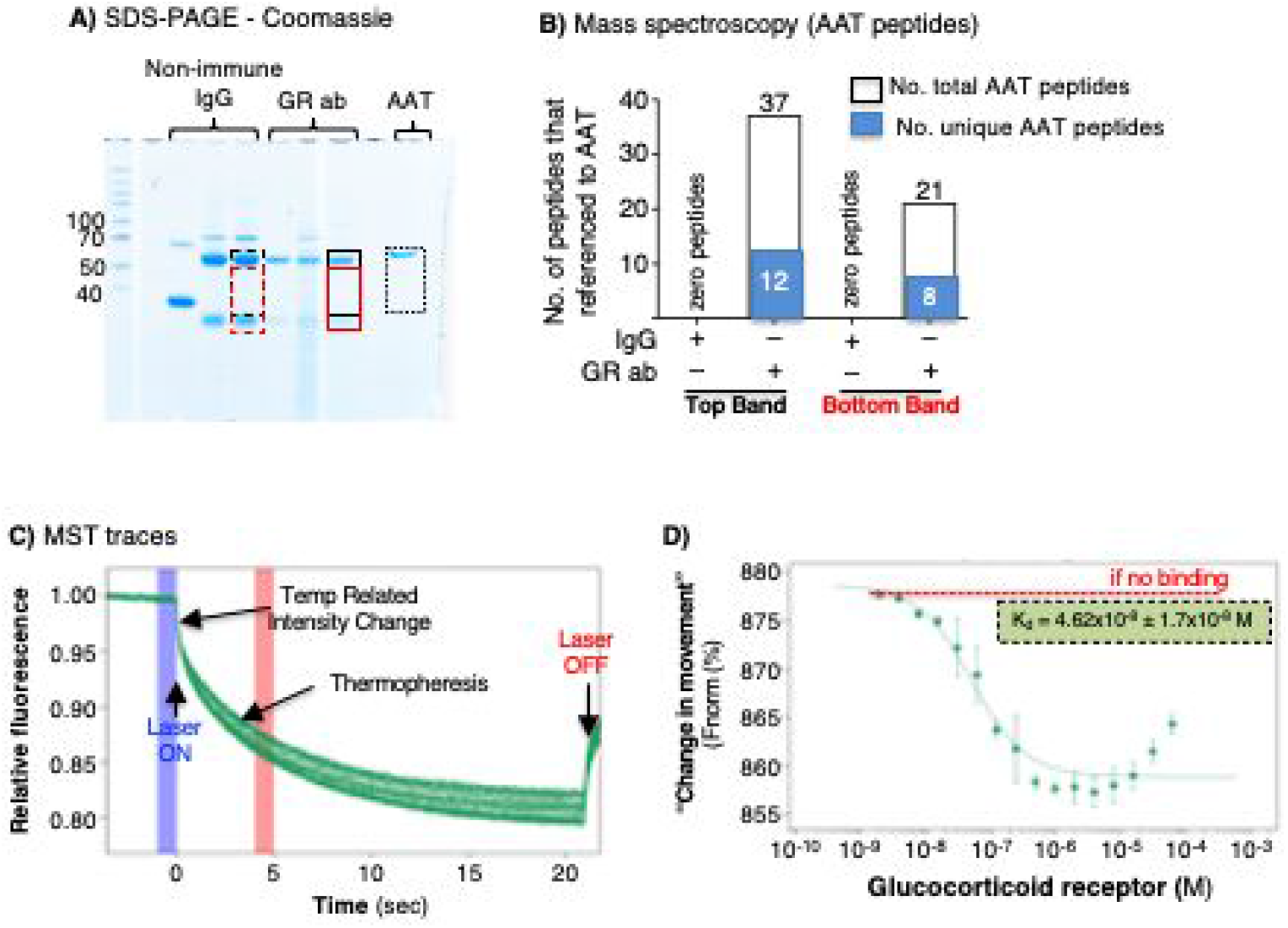
Binding of alpha-1-antitrypsin (AAT) to the glucocorticoid receptor (GR) as determined by immunoprecipitation-mass spectroscopy and in a cell-free system by microscale thermophoresis. **(A)** THP-1 macrophages were immunoprecipitated (IP’d) with anti-GR or non-immune IgG, the IP’d fractions were separated by SDS-PAGE, and the gel was stained with Coomassie blue. Stained bands in the gel (black dash and solid line boxes) that ran similarly to AAT were excised along with an area just beneath the aforementioned band (red dash and solid line boxes) and subjected to mass spectroscopy. **(B)** Graphical and numerical representation of the number of total and unique peptides that referenced to AAT in samples IP’d with control IgG (none for both the top and bottom bands) and with anti-GR (37 total and 12 unique peptides that referenced to AAT in the top band, and 21 total and 8 unique AAT peptides in the bottom band). **(C)** Microscale thermophoresis time-course tracings following the mixing of fluorescent-tagged AAT with 16 different GR concentrations and subjected to a thermogradient. **(D)** The presence of a “change in movement” of the fluorescent-tagged AAT with varying GR concentrations demonstrates that the two molecules interact *in vitro*.

For the cell lysates IP’d with anti-GR, mass spectrometry of the upper band (**Figure 3A, solid line black box**) yielded 37 total peptides that corresponded to amino acid sequences of AAT and 12 peptides that are unique to AAT (**Figure 3B**). Similarly, the lower adjacent area (**Figure 3A, solid line red box**) revealed a total of 21 peptides and 8 unique peptides that referenced to AAT (**Figure 3B**). As we had anticipated, the heavy chains of IgG also migrated to a similar position as AAT. The upper band and the adjacent lower area of the migrated lysates IP’d with anti-GR also contained 265 and 848 peptides that referenced to IgG heavy chains. However, both the upper band and lower adjacent area that were IP’d with non-immune IgG (**Figure 3A, dashed black and red boxes**, respectively) yielded zero peptides that corresponded to AAT (**Figure 3B**) but had 540 peptides and 1175 peptides that referenced to IgG. These findings confirm that AAT co-IP’d with GR.

#### Microscale Thermophoresis approach

To determine whether AAT binds to the GR in a cell-free system, we employed Microscale Thermophoresis (MST) technology. MST is a highly sensitive method that measures the mobility of fluorescently labeled proteins in the presence of a thermal gradient, a process known as thermophoresis (Jerabek-Willemsen *et al*, 2011). The thermophoretic mobility of a molecule is dependent on multiple factors, including charge, hydrodynamic radius, and interactions with the solvent. If the fluorescently labeled molecule binds another molecule, the thermophoretic mobility of the labeled molecule changes by a measurable amount. For this specific experiment, 10 nM of fluorescently labeled AAT was combined with 16 dilutions of GR ranging in concentrations from 1.91 nM to 6.25 μM. The thermophoretic mobility of AAT changed in a concentration-dependent manner with increasing concentrations of the GR in the mixture (**Figure 3C/D**). Thus, the MST studies confirmed that AAT and GR bind to each other *in vitro*.

#### Molecular modeling of GR-AAT interactions

To begin to understand how AAT and GR may interact physically, we performed *in silico* molecular modeling and docking calculations using the best two “state-of-the-art” approaches: *AlphaFold2* (Jumper *et al*, 2021) and *ClusPro* (Kozakov *et al*, 2017). GR is organized into three major domains: an intrinsically disordered N-terminal activation function-1 domain (NTD), a DNA binding domain (DBD), and a C-terminal ligand binding domain (LBD) (**Figure 4A)**. Structural modeling of the GR protein reveals the presence of two ordered domains (LBD and DBD), linked by an unstructured hinge region (**Figure 4B**). Docking revealed several plausible modes of interaction between AAT and GR both in the cytosol and the nucleus (**Figure 4**). The top docking solution from AlphaFold2 shows AAT interacting with the LBD *via* its reactive center loop (RCL), in a binding mode compatible with the GR–Hsp90/p23 cochaperone complex (Noddings *et al*, 2022) (**Figure 4C**). The AAT–GR complex is also compatible with the known dimeric arrangement of LBD (Bledsoe *et al*, 2002) (**Figure 4D**). The RCL of AAT interacts with the ligand-dependent activation function region (AF-2) in a similar fashion to nuclear coregulator proteins nuclear receptor corepressor (NCoR) and transcriptional intermediary factor-2 (TIF2; **Figure 4D**). Once translocated to the nucleus, the DBD of GR recognizes Glucocorticoid Response Elements (GREs). The top docking result using *ClusPro* positions AAT at the opposite end of the LBD, again interacting *via* its RCL, bridging the interaction with a monomeric DBD, but showing some degree of steric overlap with the second DBD formed by D-loop-mediated dimerization (**Figure 4E**).

**Figure 4.**
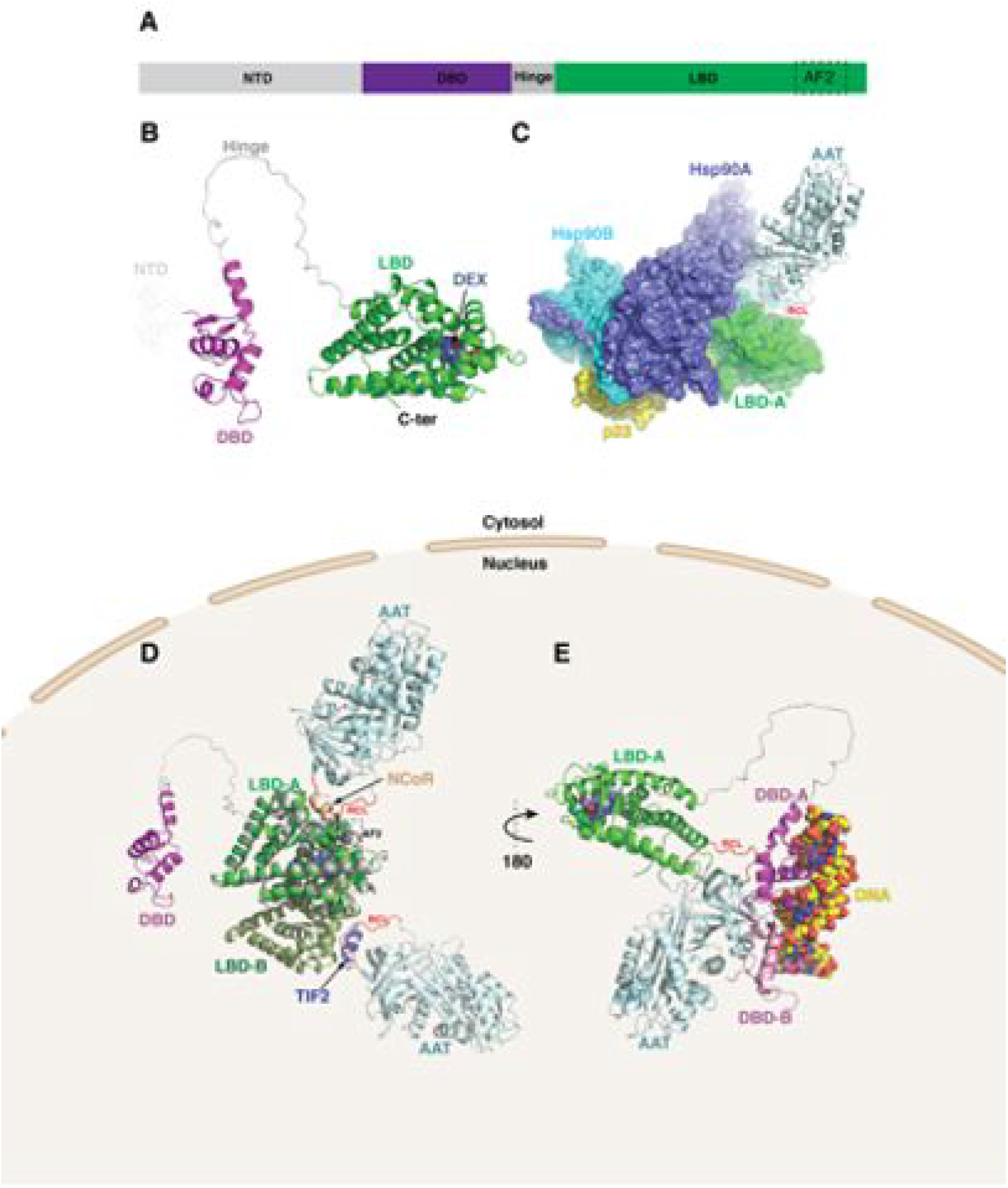
Molecular modeling of alpha-1-antitrypsin (AAT) – glucocorticoid receptor (GR) complexes. **(A)** GR is organized into three major domains: an intrinsically disordered N-terminal activation function-1 domain (NTD), a DNA binding domain (DBD), and a C-terminal ligand binding domain (LBD). In addition to its role in ligand recognition, the LBD also contains a ligand-dependent activation function region (AF-2) that is tightly regulated by hormone binding (dotted box). **(B)** Structural modeling of the GR protein reveals the presence of two ordered domains (LBD (green) and DBD (magenta)), linked by an unstructured hinge region (grey). The intrinsically disordered NTD was not modelled and is depicted by dotted lines. The C-terminus is labeled. Dexamethasone bound to the LBD is shown as steel blue spheres. **(C), (D)** and **(E)** depict three plausible modes of interactions between AAT and GR in the **(C)** cytosol and **(D, E)** nucleus. **(C)** AlphaFold2 docking simulations of LBD and AAT (PDB ID 3NE4) superimposed on the GR-Hsp90-p23 co-chaperone complex (PDB ID 7KRJ) show AAT interacting with the LBD via its RCL, in a binding mode compatible with the cochaperone complex. **(D)** The same AAT– GR complex expanded to the LBD dimer (monomers labeled LBD-A and LBD-B). Nuclear coregulator proteins nuclear receptor corepressor (NCoR; derived from PDB ID 3H52) and transcriptional intermediary factor-2 (TIF2; derived from PDB ID 1M2Z) bound to the AF-2 site (broken rectangle) are shown as brown and slate-blue cartoons, respectively. The RCL (red) of AAT interacts with the AF-2 site in a similar fashion to both nuclear regulators – involving hydrophobic interactions. Superposed (grey) is a monomeric NCoR-bound LBD (PDB ID 3H52) showing key structural differences within the AF-2 site that favor repression over activation. **(E)** In the second model of AAT binding, the RCL of AAT binds to the opposite end of LBD, potentially bridging the interaction with the DBD. The DBD is shown here in complex with DNA (space-filling atoms), modeled using the GR DNA Binding Domain monomer – TSLP nGRE Complex (magenta; PDB ID 5HN5). Also shown (pink) is a second DBD formed by D-loop-mediated dimerization (PDB ID 5E69), having some degree of steric overlap with AAT. **RCL**=reactive center loop.

### The AAT–GR complex is found in both the cytoplasmic and nuclear fractions

Given that GR can localize to the nuclear compartment, our next step in characterizing the importance of the AAT–GR interaction was to identify whether the AAT–GR complex exists in the cytoplasm, nucleus, or both. We prepared nuclear and cytoplasmic fractions of the THP-1 macrophage lysates (**Figure 5A**) and performed IP for GR, followed by AAT immunoblotting. We identified the presence of the AAT–GR complex in both the nuclear and cytoplasmic fractions, suggesting that this interaction may play a role in the transcriptional function of GR (**Figure 5B**).

**Figure 5.**
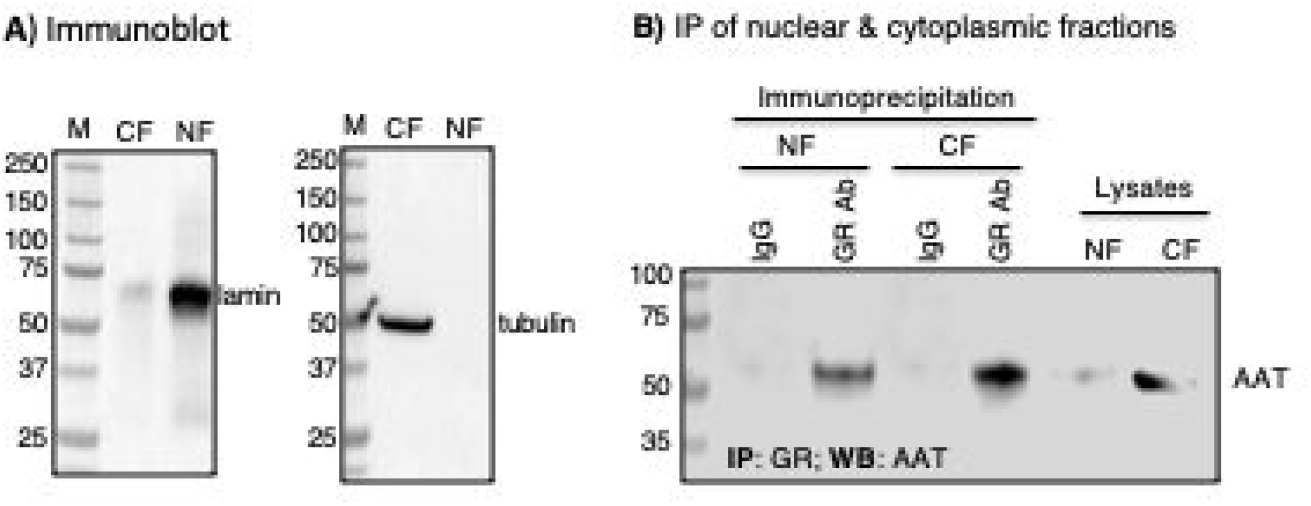
The alpha-1-antitrypsin (AAT)–glucocorticoid receptor (GR) complex is found in both the nucleus and cytoplasm. **(A)** Western blot detection of lamin and tubulin to confirm specific isolation of nuclear and cytoplasmic fractions, respectively. **(B)** Immunoprecipitation of the nuclear and cytoplasmic fractions for GR and immunoblot of immunoprecipitated lysates with an anti-AAT antibody. All data shown are representative of three independent experiments. **CF**=cytoplasmic fraction, **NF**=nuclear fraction

### Stable depletion of GR in THP-1 macrophages

We next investigated the biological importance of the AAT–GR interaction by generating a pool of transformed THP-1 macrophages where GR was stably depleted (knockdown; KD) using a lentiviral vector that expressed a shRNA-GR construct. To confirm that GR was depleted, we immunoblotted the cell lysates of THP-1^control^ and THP-1^GR-KD^ for GR, which demonstrated a significant down-regulation of GR in the THP-1^GR-KD^ cells (**Figure 6A**). To further validate the depletion of GR expression in THP-1^GR-KD^ macrophages, RNA sequencing for GR mRNA in THP-1^control^ and THP-1^GR-KD^ cells showed that THP-1^GR-KD^ had a 2.5-fold reduction in mRNA for GR compared to that of THP-1^control^ cells (**Figure 6B**).

**Figure 6.**
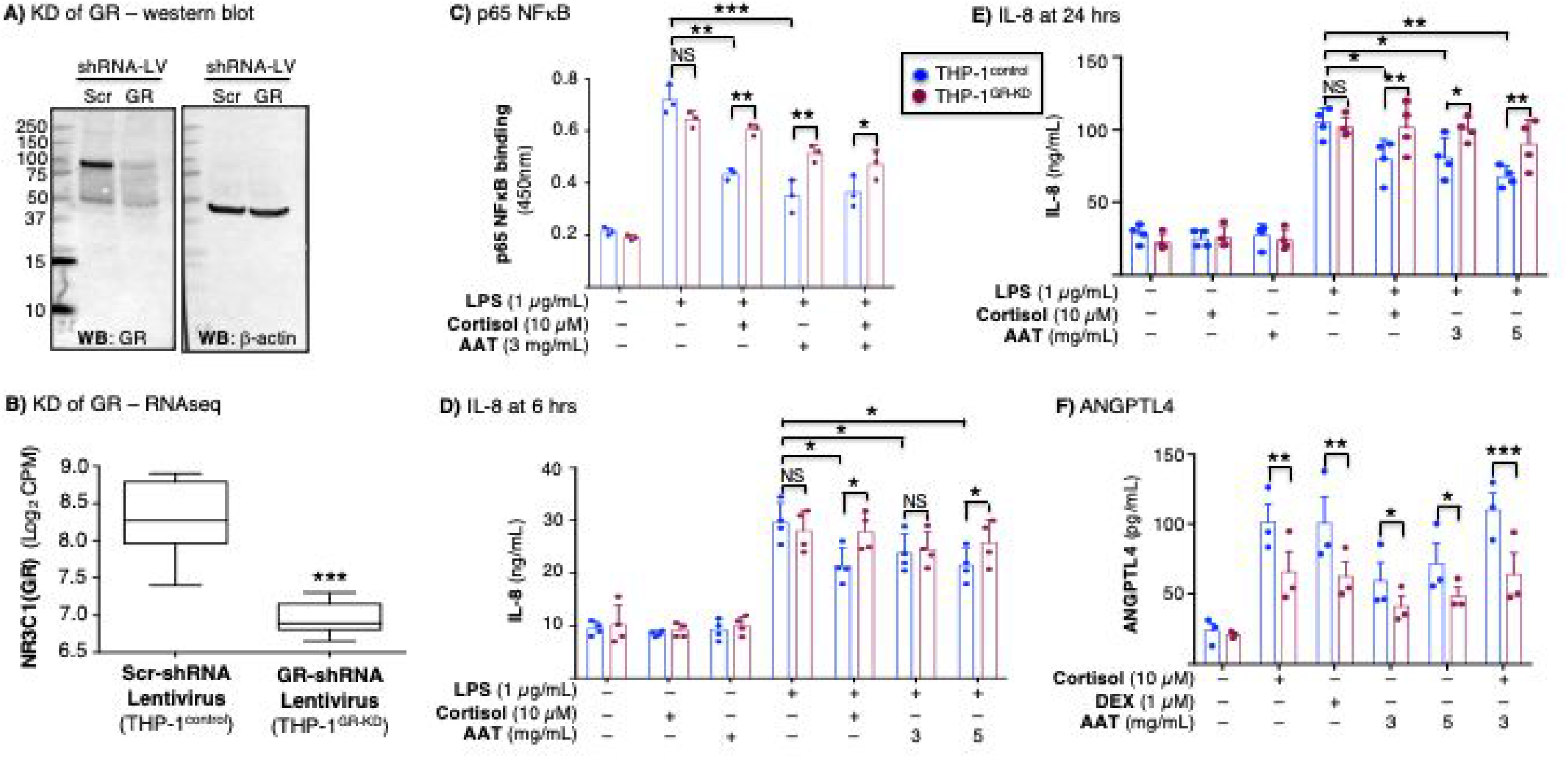
Glucocorticoids or alpha-1-antitrypsin (AAT) inhibition of NFκB activation, inhibition of interleukin-8 (IL-8) production, and induction of angiopoietin-like 4 (ANGPTL4) is glucocorticoid receptor-dependent. **(A)** Western blot of whole-cell lysates of THP-1^control^ and THP-1^GR-KD^ macrophages for GR. The data shown are representative of three independent experiments. **(B)** RNA sequencing (RNAseq) for the GR gene (*NR3C1*) transcript of the THP-1^control^ and THP-1^GR-KD^ cells using shRNA-lentivirus technology (***p<0.001 compared to Scr-shRNA lentivirus). **(C)** THP-1^control^ and THP-1^GR-KD^ macrophages were left untreated or pre-treated with cortisol, AAT, or both at the indicated concentrations for 30 minutes, and after stimulation with lipopolysaccharide (LPS) for 6 hours, p65-NFκB binding assay to its consensus oligonucleotide was performed. THP-1^control^ and THP-1^GR-KD^ macrophages were left untreated or pre-treated with cortisol, AAT, or both at the indicated concentrations for 30 minutes, and after stimulation with LPS for **(D)** 6 hours or **(E)** 24 hours, the supernatants were assayed for IL-8 by ELISA. **(F)** Glucocorticoid (cortisol or dexamethasone) or AAT induction of ANGPTL4 in THP-1^control^ and THP-1^GR-KD^ macrophages. Experiments in (C) / (F) and (D) / (E) are the mean ± SEM of three and four independent experiments, respectively, with each experiment done in duplicates. *p<0.05, **p<0.01, ***p<0.001. **Blue bars**=control shRNA (THP-1^control^), **red bars**=GR shRNA (THP-1^GR-KD^). **NS**=not significant, **KD**=knockdown, **LV**=lentivirus, **Scr**=scrambled.

### The AAT–GR complex inhibits lipopolysaccharide-induced NFκB activation and IL-8 production

Since GCs have potent anti-inflammatory function *via* the ability of the GC–GR complex to affect inflammatory gene transcription related to NFκB targets (Necela & Cidlowski, 2004; Van Bogaert *et al*, 2010), we examined the role of AAT in NFκB activation in the context of GR signaling. We pre-treated THP-1^control^ and THP-1^GR-KD^ macrophages with cortisol, AAT, or both for 30 minutes, then stimulated the cells with lipopolysaccharide (LPS) for 6 hours, and quantified binding of p65-NFκB to its consensus oligonucleotide. In THP-1^control^ cells, LPS induced p65-NFκB binding was significantly inhibited by cortisol or AAT (**Figure 6C, blue bars**). However, although LPS-induced activation of p65-NFκB in THP-1^GR-KD^ macrophages was comparable to THP-1^control^ cells (**Figure 6C, second set of red and blue bars**, respectively), there was less inhibition by cortisol or AAT in the THP-1^GR-KD^ macrophages (**Figure 6C, compare the second set of bars to their corresponding last three sets of bars**). Furthermore, whereas the inhibitory effect of cortisol was nearly completely abrogated in the THP-1^GR-KD^ macrophages (**Figure 6C, second vs. third red bars**), the inhibitory effects of AAT were not completely attenuated in the THP-1^GR-KD^ macrophages (**Figure 6C, second vs. fourth and fifth red bars**), suggesting that AAT also affects NFκB signaling in a GR-independent manner.

To compare LPS-induced expression of IL-8 in the supernatants of THP-1^control^ and THP-1^GR-KD^ macrophage cultures, both cell types were stimulated with LPS for 6 and 24 hours alone or in the presence of cortisol (10 μM) or AAT (3 and 5 mg/mL), and the supernatants measured for IL-8 expression. The normal AAT level in plasma is 1-2 mg/mL but may increase 3- to 5-fold in states of systemic inflammation and/or infection (de Serres *et al*, 2003; Hazari *et al*, 2017). There was robust induction of IL-8 by LPS alone at 6 and 24 hours in both THP-1^control^ and THP-1^GR-KD^ macrophages, with greater amounts accumulated at the longer time point (**Figures 6D/E**). In THP-1^control^ macrophages, there was significant inhibition of LPS-induced IL-8 expression following incubation with cortisol or 5 mg/mL AAT, particularly at 24 hours (**Figures 6D/E, blue bars**). However, in THP-1^GR-KD^ macrophages, neither cortisol nor AAT significantly inhibited LPS-induced IL-8 production (**Figures 6D/E, red bars**).

### AAT induction of angiopoietin-like 4 protein is mediated by GR

It has been established that AAT or GCs have the ability to induce the expression and release of angiopoietin-like 4 protein (ANGPTL4) in human blood monocytes and human microvascular endothelial cells (Frenzel *et al*, 2015; Koliwad *et al*, 2009; Nakamoto *et al*, 2017). Thus, THP-1^control^ and THP-1^GR-KD^ macrophages were stimulated with cortisol (10 μM), dexamethasone (1 μM), AAT (3 and 5 mg/mL), or both cortisol and AAT for 24 hours and the supernatants were used to quantify the ANGPTL4 levels (Yin *et al*, 2009). Stimulation of THP-1^control^ macrophages with cortisol, dexamethasone, or AAT induced production of ANGPTL4, although stimulation with AAT was not as robust as the GCs (**Figure 6F, blue bars**). However, GC- or AAT-induced expression of ANGPTL4 was significantly lower in the THP-1^GR-KD^ macrophages (**Figure 6F, red bars**).

### AAT–GR binding enhances mycobacterial control in THP-1 macrophages

The consensus among clinicians and scientists is that GCs increase susceptibility to *Mycobacterium tuberculosis* (Jick *et al*, 2006; Lai *et al*, 2015) and possibly non-tuberculous mycobacteria (NTM) (Xie *et al*, 2021). On the other hand, AAT enhances autophagy, an effector mechanism known to kill intracellular mycobacteria (Bai *et al.*, 2019; Bai *et al*, 2013; Deretic *et al*, 2006; Gutierrez *et al*, 2004). Since canonical GR signaling and the cellular effector function(s) induced by AAT appear to have opposing effects in controlling mycobacterial infection, we examined in macrophages the role of the AAT–GR interaction in the control of *M. tuberculosis* infection and the NTM, *Mycobacterium intracellulare*. THP-1^control^ and THP-1^GR-KD^ macrophages were initially infected with *M. tuberculosis* H37Rv or *M. intracellulare* at a multiplicity-of-infection of 10 mycobacteria:1 macrophage for 1 hour, 2 days, and 4 days (without exogenous GC or AAT added), and then the mycobacteria were quantified. We detected a productive *M. tuberculosis* and *M. intracellulare* infection of the THP-1^control^ macrophages, peaking at day 2 (**Figure 7A, top and bottom panels, blue bars**), and which, unexpectedly, consistently increased in the THP-1^GR-KD^ macrophages (**Figure 7A, top and bottom panels, red bars**).

**Figure 7.**
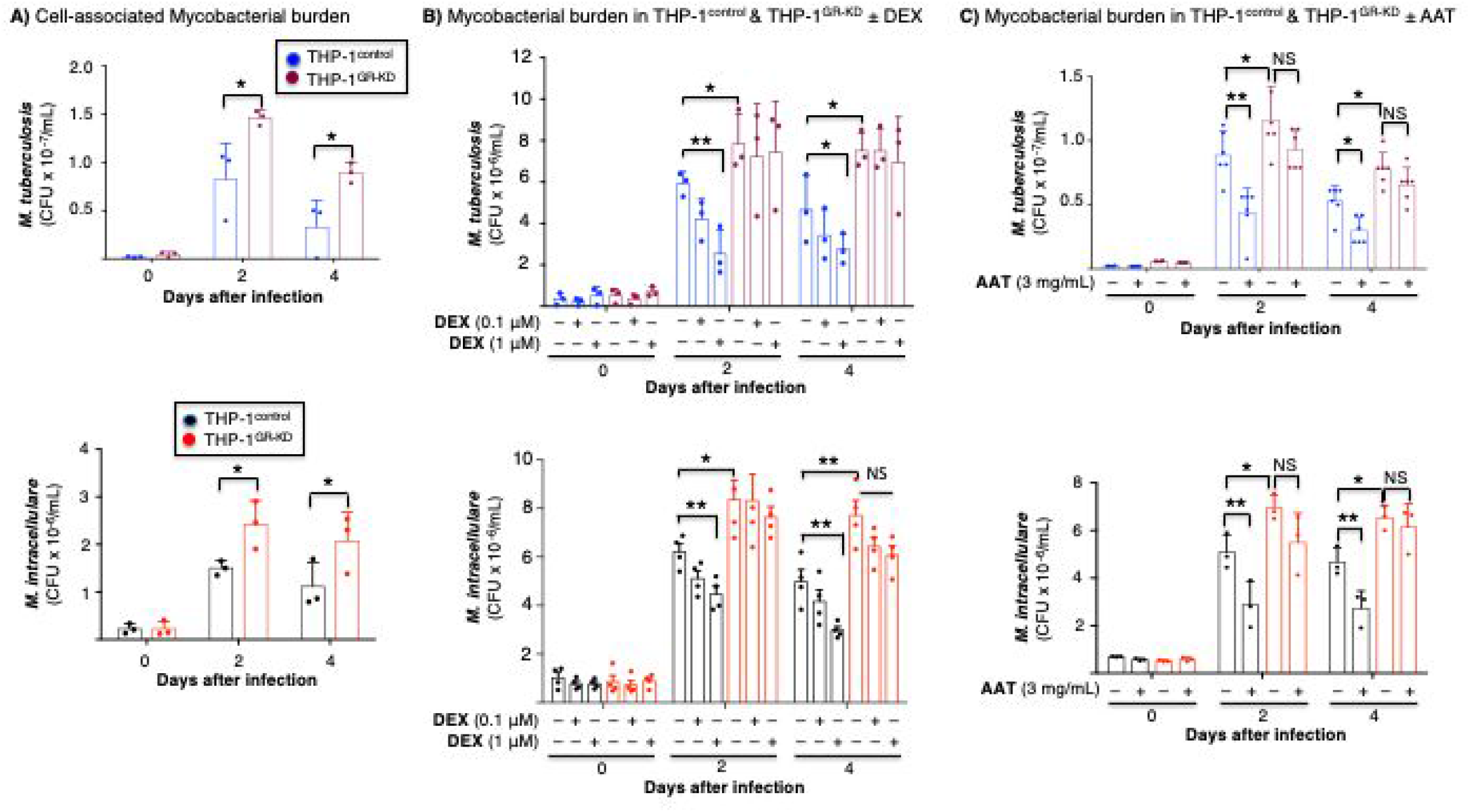
Mycobacterial infection of THP-1^control^ and THP-1^GR-KD^ macrophages in the absence or presence of alpha-1-antitrypsin (AAT) or glucocorticoid. **(A)** THP-1^control^ and THP-1^GR-KD^ macrophages were infected with *M. tuberculosis* H37Rv or *M. intracellulare* for 1 hour, 2 and 4 days. The cells were washed, and intracellular mycobacteria quantified. **(B)** THP-1^control^ and THP-1^GR-KD^ macrophages were left untreated or pre-treated with dexamethasone (0.1 or 1 μM) for 60 minutes, followed by *M. tuberculosis* or *M. intracellulare* infection for 1 hour, 2 days, and 4 days. The cells were washed and intracellular mycobacteria quantified. **(C)** THP-1^control^ and THP-1^GR-KD^ macrophages were left untreated or pre-treated with 3 mg/mL AAT for 30 minutes, infected with *M. tuberculosis* or *M. intracellulare* for 1 hour, 2 and 4 days, the cells washed, and intracellular mycobacteria quantified. Experiments in (A) and (B) are the mean ± SEM of three independent experiments, and in (C), the mean ± SEM of six independent experiments, with each experiment done in duplicates. *p<0.05, **p<0.01. **Blue bars**=control shRNA (THP-1^control^), **red bars**=GR shRNA (THP-1^GR-KD^). **NS**=not significant.

Since depletion of basal GR increased burden of both mycobacterial species in THP-1 macrophages, we sought to determine the effects of GCs in the control of *M. tuberculosis* and *M. intracellulare*. THP-1^control^ and THP-1^GR-KD^ macrophages were pre-incubated with 0.1 and 1 μM dexamethasone for 60 minutes and then infected with *M. tuberculosis* or *M. intracellulare*. One hour, 2 days, and 4 days after infection, the infected THP-1 cells were washed, lysed, and each of the mycobacterial species quantified by counting CFUs on solid agar plates. In THP-1^control^ macrophages, dexamethasone *reduced* the burden of both *M. tuberculosis* and *M. intracellulare* in a dose-dependent fashion (**Figure 7B, top and bottom panels, blue bars**). However, dexamethasone had no significant effect on the number of the mycobacterial CFUs in the THP-1^GR-KD^ macrophages (**Figure 7B, top and bottom panels, red bars**).

To determine the effects of AAT on *M. tuberculosis* and *M. intracellulare* infection in the context of GR, THP-1 ^control^ and THP-1^GR-KD^ macrophages were then pre-incubated with AAT (3 mg/mL) for 30 min, infected with *M. tuberculosis* or *M. intracellulare*, and mycobacteria quantified after 1 hour, 2 days, and 4 days after infection. In THP-1 ^control^ macrophages, preincubation with AAT significantly reduced the burden of both *M. tuberculosis* and *M. intracellulare* (**Figure 7C, top and bottom panels, blue bars**), whereas this did not occur in THP-1^GR-KD^ macrophages (**Figure 7C, top and bottom panels, red bars**). While the ability of dexamethasone to enhance macrophage control of *M. tuberculosis* (or *M. intracellulare*) is not consistent with the paradigm that GC use may predispose patients to tuberculosis (TB) and NTM infections, other studies have shown that GCs can upregulate a host-protective innate immune response through increased expression of Toll-like receptor (TLR)2 and enhanced TLR4 signaling (Busillo & Cidlowski, 2013; Chinenov & Rogatsky, 2007; Zhang & Daynes, 2007). Thus, to determine whether GC alone can induce TLR2, a pattern recognition receptor capable of recognizing lipoproteins of mycobacteria, we immunoblotted for TLR2 in unstimulated cells or with dexamethasone stimulation in THP-1^control^ and THP-1^GR-KD^ macrophages. Notably, dexamethasone induced TLR2 expression in the THP-1 ^control^ macrophages, but its expression was markedly diminished in the THP-1^GR-KD^ macrophages (**Figure 8**).

**Figure 8.**
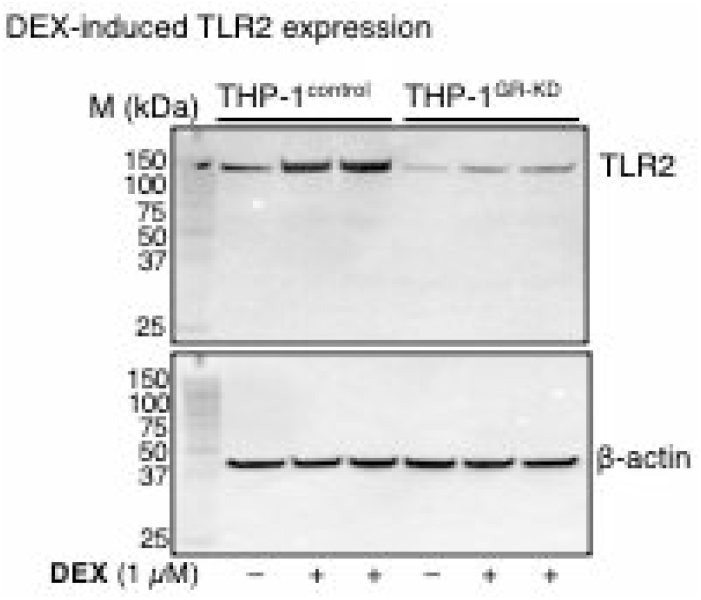
Glucocorticoid induces expression of a pattern-recognition receptor. THP-1^control^ and THP-1^GR-KD^ macrophages were left untreated or pre-treated with dexamethasone (1 μM) for 24 hours, followed by immunoblotting of the whole-cell lysates for TLR2 (top). The same membrane was stripped and immunblotted for β-actin (bottom). Data shown in representative of three independent experiments.

## DISCUSSION

Herein we demonstrated by several lines of evidence that AAT binds directly to the GR in primary human MDMs and/or a macrophage cell line, including tandem IP-immunoblotting, IP-mass spectrometry, and a cell-free system using microscale thermophoresis. We further showed in THP-1^GR-KD^ macrophages that AAT–GR binding has biological consequences. Specifically, AAT inhibited LPS-induced NFκB activation and IL-8 production, induced ANGPTL4 expression, and reduced *M. tuberculosis* burden in macrophages in a GR-dependent fashion. However, alternative pathways, such as Akt signaling, may contribute to inhibiting both LPS-induced NFκB activation and IL-8 production by GR signaling in this system (Hattori *et al*). But the inhibitory effects of cortisol on NFκB activation were more pronounced than the inhibitory effects of AAT in the THP-1^GR-KD^ macrophages (Figure 5A), suggesting that AAT affects signaling and gene regulation in both GR-dependent and GR-independent fashion (Chan *et al*, 2012; Shapiro *et al.*, 2001).

Individuals with AAT deficiency have increased neutrophilic pulmonary infiltration and proclivity to neutrophil-mediated panniculitis or vasculitis due to the ability of AAT to inhibit neutrophil infiltration by two reported mechanisms: *(i)* the ability of the glycan moieties of AAT to bind IL-8, preventing the neutrophil chemokine from binding to its receptor (CXCR1) and *(ii)* the ability of AAT to inhibit ADAM17, a disintegrin capable of cleaving cell membrane-bound FcγRIIIb bound to immune complexes which then binds complement receptor 3 to induce cytoskeletal rearrangements necessary for neutrophil chemotaxis (Bergin *et al.*, 2010; McCarthy *et al*, 2016). We now demonstrate a third mechanism where AAT promotes GR-mediated inhibition of IL-8 production *via* antagonizing proinflammatory NFκB activation.

Ultimately, while we have established the AAT–GR physical interaction and the physiologic role for this interaction in antagonizing NFκB-mediated inflammation and promoting the transcription of target genes (**Figure 9A**), the molecular mechanism by which AAT binds GR and regulates transcription remains to be defined. While AAT is very unlikely to act as a GR ligand through the canonical GR ligand binding domain (as supported by molecular modeling), several alternative avenues that are not necessarily mutually exclusive may explain its role in GR function, forming the basis for further investigations (**Figure 9B**). First, our IP findings of the nuclear and cytoplasmic fractions suggest that GR shuttles between the two compartments (**Figure 9B1**). AAT binding to GR may increase GR localization to the nuclear compartment compared to absent or lower AAT levels. Second, exogenous or supraphysiologic AAT may act primarily in the cytoplasm to modulate the disassembly of GR from chaperone proteins and facilitate nuclear translocation and transactivation upon binding to canonical ligands (**Figure 9B2**). We observed GR-dependent inhibition of NFκB activity in response to AAT but in the absence of exogenous GC. This may be due to low concentrations of GC in the cell culture conditions potentiated by AAT. A less likely possibility is that of a direct transcriptional role for AAT that facilitates translocation of GR to the nucleus where GR activates transcription in a ligand-independent manner although each is not necessarily mutually exclusive of the other. A third possibility of the cooperation between AAT and the GC–GR is that the serine protease inhibitor activity of AAT may exert an indirect, stabilizing effects of some transcriptional complexes that interact with GR, which may occur via its serine protease inhibitory activity or by a yet uncharacterized mechanism (**Figure 9B3**).

**Figure 9.**
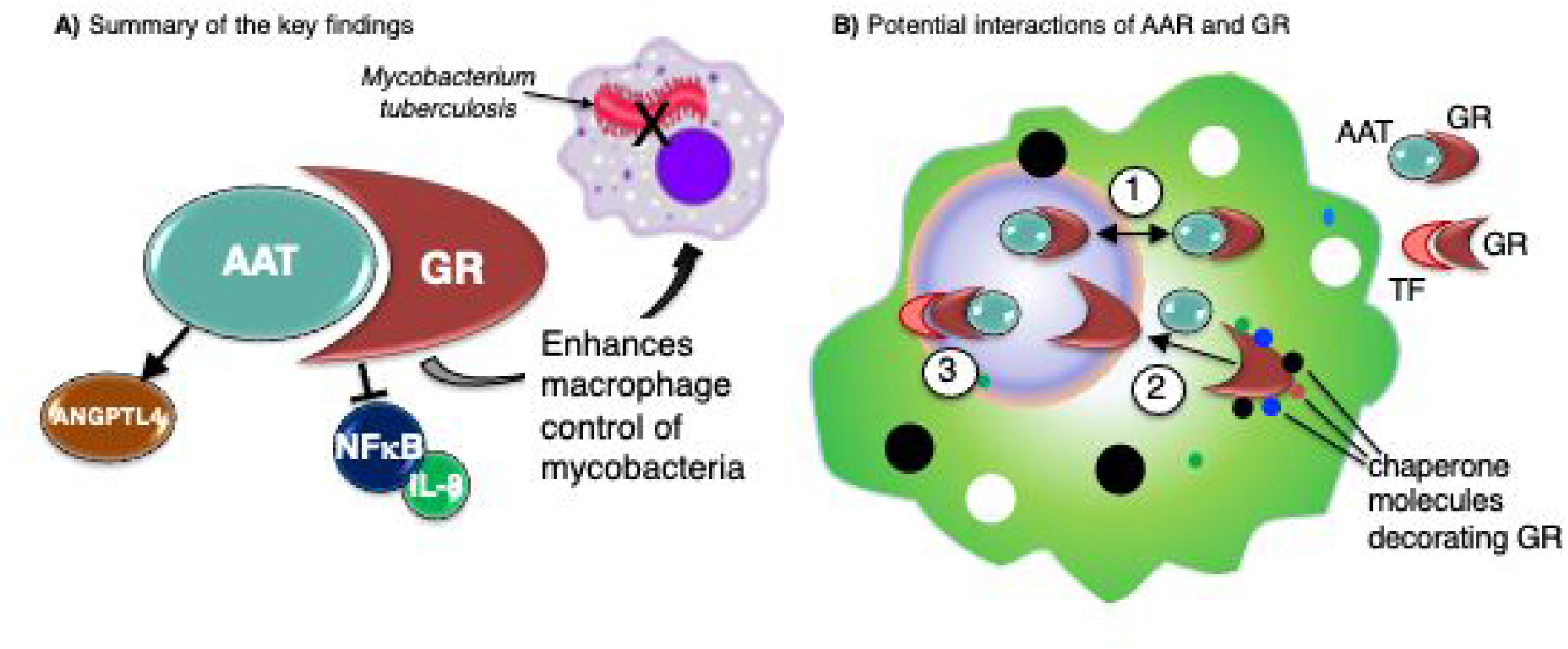
Illustration of the key findings and hypotheses of the AAT–GR interactions. **(A)** In summary of the key findings, alpha-1-antitrypsin-glucocorticoid receptor (AAT–GR) interaction inhibits nuclear factor-kappa B (NFκB) activation and interleukin-8 (IL-8) production, induces angiopoietin-like 4 (ANGPTL4), and enhances macrophage control of *Mycobacterium tuberculosis*. **(B)** Several hypotheses of how AAT–GR interaction may affect cellular function: (1) AAT may shuttle GR between the nuclear and cytoplasmic compartments; (2) AAT may facilitate disassembly of GR–chaperone complexes; (3) AAT may stabilize transcriptional complexes of GR–transcription factor (orange structure) or locally modulate the activity of proteases.

We performed molecular modeling of AAT–GR complexes to explore these possibilities further. AAT binding to the LBD of GR is compatible with cochaperone-GR interactions in the cytosol (Figure 4C), suggesting that AAT may play a role in assembly/disassembly of this complex. Intriguingly, this complex positions the AAT RCL interacting with the AF-2 region in a similar mode to nuclear coregulator proteins NCoR and TIF2 (Figure 4D). Although the RCL does not contain the conserved Lxxx(I/ L)xxx(I/L) motif found in corepressors (Frank *et al*, 2021), its hydrophobic nature and flexibility is compatible with the hydrophobic AF-2 site. Conformational dynamics of the AF-2 site is known to dictate its function as a coactivator or a corepressor site (Köhler *et al*, 2020; Schoch *et al*, 2010), and is apparent from structural comparisons of LBD-ligand complexes (Figure 4D). Coregulator molecules binding to AF-2 tune its conformational dynamics, as well as its cooperativity with the GC ligand binding site, and thus govern downstream events; for example, release of bound chaperones, nuclear translocation, dimerization, DNA-binding, and ultimately transcription. It is therefore plausible that AAT binding to the AF-2 site may similarly achieve allosteric regulation of downstream activation and repression pathways. An alternative docking solution positions AAT at the opposite end of the LBD, again interacting *via* its RCL and bridging the interaction with a monomeric DBD; however, this model showed some degree of steric overlap with the second DBD formed by D-loop-mediated dimerization (Figure 4E). This mode of interaction within the nucleus prompts speculation that AAT binding may favor monomer over dimer and thus the repression of pro-inflammatory genes (Hudson *et al*, 2013). However, our finding that AAT induces ANGPTL4 expression, together with the known dimeric GR DBD that binds to the ANGPTL4 GRE (Koliwad *et al.*, 2009), is more consistent with a scenario where AAT is released upon dimerization, which can then exert localized effects, be it protease-inhibition or other binding activity that may be key for gene induction. However, the flexible nature of the hinge region limits the accuracy of the modeling and therefore such interpretations must be made with caution. Nevertheless, overall, modeling highlights several experimentally-testable hypotheses on the role of AAT in GR biology.

Our findings suggest that short-term GR activation (whether by GC or AAT) in differentiated THP-1 macrophages directs protective responses against mycobacteria (Figure 9A). On the surface, this notion is counterintuitive given the well-described pharmacologic role of GC in suppressing immunity, resulting in predisposition to TB (Jick *et al.*, 2006; Lai *et al.*, 2015). However, it is important to emphasize that GC-induced susceptibility to TB occurs with GC doses that are significantly greater than physiologic output and for an “extended period” of time *in vivo*. Yet, we demonstrate in two lines of evidence that GR activation enhances THP-1 macrophage control of *M. tuberculosis* and *M. intracellulare: (i)* the downregulation of GR in the absence of exogenous AAT (or GC) resulting in increased *M. tuberculosis* and *M. intracellulare* burden in the differentiated mononuclear cells and *(ii)* the treatment of THP-1 macrophages with either AAT or GC reduced the burden of *M. tuberculosis* and *M. intracellulare* in a GR-dependent fashion. This seemingly paradoxical finding is supported by the literature. Tukenmez *et al* (Tükenmez *et al*, 2019) found that many different GC formulations individually decreased *M. tuberculosis* burden in human MDMs. Bongiovanni and co-workers (Bongiovanni *et al*, 2015) similarly reported that cortisol not only decreased phagocytosis of *M. tuberculosis* (by ~10%) but that there was a further reduction in the intracellular burden of *M. tuberculosis* by day 4 of culture in THP-1 macrophages. In contrast, Wang *et al* (Wang *et al*, 2017) found that dexamethasone enhanced the growth of *M. bovis* BCG in primary mouse and human macrophages, raising the possibility that the effects of GC on macrophage control of mycobacterial infection may be mycobacterial species- and host-specific. It cannot be overemphasized that such potential host-protective effects of GCs in pure macrophage cultures are likely to be different *in vivo*, where, for example, GCs in pharmacologic doses induce apoptosis in lymphocytes, which are critical in controlling mycobacterial infection. Thus, future elucidation on how GC may be exploited to maximize innate immune function while minimizing their immunosuppressive effects on lymphocytes may bring new approaches to treat mycobacterial infections.

In support of the aforementioned findings, other studies have shown that low-dose GC has immunoenhancing effects in myeloid cells (Barber *et al*, 1993; Busillo & Cidlowski, 2013; Fantuzzi *et al*, 1995; Smyth *et al*, 2004; Zhang & Daynes, 2007). GCs have been shown to enhance immunity by inducing TLR2 expression, especially in the presence of TNF and IL-1β (Busillo & Cidlowski, 2013; Chinenov & Rogatsky, 2007). We also showed that dexamethasone induced TLR2 expression in a GR-dependent fashion (Figure 8). TLR2 activation may also induce autophagy (Seto *et al*, 2012), and various polymorphisms of TLR2 have been associated with an increased risk for TB and NTM lung disease (Bento *et al*, 2015; Yim *et al*, 2008). Zhang and Daynes (Zhang & Daynes, 2007) demonstrated that during differentiation of murine myeloid progenitors into macrophages, incubation with corticosterone increased the cellular responsiveness to LPS through increased expression of a phosphatase that inhibits PI3 kinase/Akt with subsequent increase in TLR4 signaling. Since we found that GC inhibited LPS-induction of NFκB activation and IL-8 production, possible explanations that may account for these seemingly contradictory findings are: *(i)* species differences in the phenotypic response to GC, *(ii)* differential responses to singular GC formulations, *(iii)* different signaling pathways that may oppose each other may be similarly affected by GC, and *(iv)* presence of a bimodal dose response paradigm wherein lower GC doses are immunopermissive whereas higher doses are immunosuppressive (Zhang & Daynes, 2007). Our previous findings that inhibition of NFκB activation in macrophages (by AAT or a small molecule inhibitor of IκBα) enhanced autophagy and clearance of *M. intracellulare* or *M. tuberculosis* infections would lend further credence that GC inhibition of NFκB activation augments macrophage activity against mycobacteria (Bai *et al.*, 2019; Bai *et al.*, 2013).

We previously showed that AAT induces autophagosome formation and maturation in mycobacteria-infected macrophages, which enhances bacterial clearance (Bai *et al*., 2019). But could AAT have effects in other cell types that are beneficial in the context of TB? One possibility is that AAT antagonizes the damaging effects of neutrophils in the later stages of TB (Lowe *et al*, 2012) by: *(i)* sequestering IL-8 (Bergin *et al.*, 2010), inhibiting neutrophil chemotaxis (McCarthy *et al*., 2016), and, as we have shown, inhibiting IL-8 production *via* binding to GR and *(ii)* inhibiting formation of neutrophil extracellular traps (NETs) – which are elastase- and IL-8-induced extruded DNA decorated with citrullinated histones, elastase, and other proteins – meant to capture extracellular bacteria but may also contribute to tissue injury (de Melo *et al*, 2018; Dömer *et al*, 2021; Frenzel *et al*, 2012; Wartha *et al*, 2007). AAT has also been shown to cause decreased shedding of macrophage mannose receptors (MMR), increasing their cell surface expression and reducing soluble MMR, effects which may contribute to the antiinflammatory effects of AAT (Embgenbroich *et al*, 2021; Serban *et al*, 2017).

A limitation of this study is that we used a differentiated monocytic cell line to knock-down GR to study the biological significance of the AAT-GR interaction. On the other hand, the advantage of using THP-1 cells to create a pool of immortal cells stably knocked down for GR is that transient knockdown of primary macrophages would introduce more confounding variables such as non-clonality of cells, differential epigenetics among different individuals, the influence of AAT variants and glucocorticoid resistance (latter often due to genotypic differences in GR subunits), and the acute cellular stress associated with transient transfection to accomplish gene knockdown. Nevertheless, future studies using primary macrophages and accounting for the aforementioned variables to determine the biological significance of the AAT-GR interaction are likely to lead to additional mechanistic insights to this paradigm. Another limitation is that we used a laboratory strain of *M. tuberculosis* as clinical strains of varying virulence may have more robust immune evasive mechanisms. While subjects with AAT deficiency are predisposed to NTM lung disease, perhaps a third limitation is that we are unaware such individuals are also predisposed to TB as our findings may imply. Possible reasons for this lattermost point are that AAT anomalies are uncommon in most of the TB endemic areas of the world due mostly to racial differences (*i.e*., clinically significant AAT variants are more common in the Caucasian population), AAT phenotype or levels are not checked in the vast majority of cases of TB, and because *M. tuberculosis* is relatively more virulent than NTM and can affect healthy individuals, the number of TB cases that may be associated with AAT deficiency will be a small minority of the entire TB population.

In conclusion, AAT is a highly abundant biomolecule with a well-demonstrated protective role against multiple inflammatory disorders and disease states. The AAT–GR interaction we described intersects two crucial anti-inflammatory pathways that are highly amenable to pharmacologic manipulation. We have shown that AAT–GR interactions can modulate inflammation and infection with *M. tuberculosis*. A novel implication of AAT–GR biology is an alternative or adjunct to GC therapy in treating chronic inflammation due to non-infectious or infectious causes. Indeed, comparable induction of GR responses between exogenous AAT administration and GCs opens the door for novel methods to manipulate one of the most potent and widely prescribed anti-inflammatory pharmacologic targets.

## EXPERIMENTAL PROCEDURES

### Materials

The human monocytic cell line (THP-1) and *Mycobacterium tuberculosis* H37Rv were obtained from the ATCC (Manassas, Virginia). *M. intracellulare* NJH9141 is a clinical strain isolated at National Jewish Health. Fetal bovine serum (FBS) was purchased from Atlanta Biologicals (Lawrenceville, GA) and heat-inactivated at 56°C for 30 minutes. Recombinant human GR protein (#A15663), GR-specific polyclonal antibody (rabbit) (PA1-551A) with robust GR specificity in applications such as GR chromatin IP (John *et al*, 2011), AAT-specific polyclonal antibody (rabbit), AAT-specific monoclonal antibody (2B12) (mouse), and the human angiopoietin-like 4 protein (ANGPTL4) ELISA Kit were purchased from Invitrogen-ThermoFisher Scientific (Carlsbad, CA). TLR2-specific polyclonal antibody (rabbit) (#12276), tubulin polyclonal antibody (#2144), lamin B polyclonal antibody (#12586), and β-actin polyclonal antibody (rabbit) were purchased from Cell Signaling Technology (Danvers, MA). Human IL-8/CXCL8 Quantikine ELISA kit and non-immune human IgG control were purchased from R&D Systems (Minneapolis, MN). Phorbol myristate acetate (PMA), lipopolysaccharide (#L-2880), cortisol (#C-106), and dexamethasone (#1756) were purchased from Sigma (St. Louis, MO). The TransAM^®^ NFκB p65 kit was purchased from Active Motif (Carlsbad, CA). Glucocorticoid Receptor (NR3C1) human shRNA (shRNA-GR) Lentiviral Particles were a kind gift from Miles Pufall, Ph.D. (University of Iowa Carver College of Medicine). Protein A Sepharose™ 4 Fast Flow sepharose beads were purchased from GE Healthcare Life Sciences (Pittsburgh, PA). AAT (Glassia^®^) was acquired from Kamada Ltd., Israel.

### Macrophage culture

Human THP-1 cells were cultured in RPMI-1640 medium containing 2 mM L-glutamine (Gibco; Grand Island, NY), 10% FBS, penicillin (100 U/mL), and streptomycin (100 μg/mL) at 37°C and 5% CO_2_ and in the presence of 15 ng/mL PMA for 24 hours to induce differentiation into macrophages. Murine RAW 264.7 cells were cultured in the same medium but without PMA. Peripheral blood monocytes from a healthy adult donor were isolated from blood collected in CPT^TM^ tubes after informed consent based on an approved Institutional Review Board protocol (HS-2651) that abides by the Declaration of Helsinki principles. Differentiation of the human monocytes to monocyte-derived macrophages (MDM) was performed in the presence of 20 ng/mL monocyte-colony stimulating factor and cultured in 10% autologous plasma as previously described (Bai *et al.*, 2019).

### Fluorescent immunocytochemistry

Differentiated human THP-1 macrophages and MDMs, and the murine RAW 246.7 monocytic cell line were seeded onto 4-well chamber slides (Lab-Tek; ThermoFisher Scientific Inc.) at a density of 0.5-1 x10^5^ cells/well. Cells were fixed with 4% paraformaldehyde for 30 minutes. Fixed cells were permeabilized for 20 minutes using 0.3% Triton X-100 in PBS. Nonspecific binding of antibodies was blocked by incubating the cells for 1-1.5 hours 3% BSA dissolved in 1X PBST. Cells were incubated with polyclonal anti-GR (1:300 dilution) and monoclonal anti-AAT (1:300 dilution) antibodies overnight at 4°C, washed thrice, and incubated with anti-rabbit CY3 and anti-mouse Alexa Fluor secondary antibodies (both 1:1000 dilution) at room temperature for 1-2 hours. The slides were sealed with ProLong Gold^™^ Antifade Mountant with DAPI (ThermoFisher Scientific). The THP-1 macrophages were imaged using a SP5 confocal microscope (Leica Microsystems) with a magnification of 63X and a 1.4 numerical aperture (NA) oil immersion objective. The fluorescent labels, DAPI, anti-AAT, and anti-GR were excited using 405 nm, 488 nm, and 543 nm laser excitation, respectively. The MDM and RAW 264.7 macrophages were analyzed at 40X magnification using a fluorescent microscope (Carl Zeiss Axiovert 200M) equipped with DAPI and fluorescent filters.

### Immunoprecipitation (IP) and immunoblot analyses

IP was performed according to a previously published protocol (Bai *et al*, 2009). Briefly, the cells were lysed with 50 mM Tris–HCl, pH 7.4, containing 0.5% (v/v) Nonidet P-40, 1 mM EDTA, 150 mM NaCl, 5 μg/ml aprotinin, 5 μg/ml leupeptin, 2 mM Na3VO4, and 1 mM PMSF. Then 300 μg of protein from whole cell lysate preparations were incubated with 3 μg of anti-GR polyclonal antibodies, anti-AAT polyclonal antibodies, or control non-immune IgG at 4°C overnight with gentle rotary mixing. The protein-antibody complexes were isolated by incubation with 20 μL protein A-sepharose beads with gentle mixing for 2 hours at 4°C. The beads were then washed three times with wash buffer (10 mM Tris buffer, pH 7.4 with protease inhibitor cocktail). The bound immune complexes were eluted by heating in loading buffer and analyzed for AAT or GR by western blot.

Each immunoprecipitated (IP’d) pellet was resuspended in 30 μL 1X Laemmli/DDT buffers and heated at 95°C for 5 minutes for immunoblot analysis. Either the IP’d preparations or whole-cell lysates were fractionated using 12% SDS-PAGE and transferred onto PVDF membranes using the iBlot 2 dry blotting System from Invitrogen (ThermoFisher Scientific). For immunodetection of AAT, the membranes were blocked in PBST buffer containing 5% fat-free milk powder (blocking buffer) for one hour, then incubated overnight at 4°C with anti-AAT polyclonal antibody, anti-AAT monoclonal antibody, or anti-β-actin antibody (each at a dilution of 1:1000 (v/v)), followed by detection with HRP-conjugated anti-rabbit IgG (1:2000 dilution) or HRP-conjugated anti-mouse IgG (1:2000). To detect GR, polyclonal anti-GR polyclonal (1:1000) or anti-β-actin (1:1000) were diluted in PBST buffer with 5% BSA overnight at 4°C with light shaking, followed by detection using HRP-conjugated anti-rabbit IgG at 1:2000 to detect GR or 1:5000 to detect ß-actin-antibody binding in blocking buffer at room temperature for 1-2 hours. Immunoblotting for lamin and tubulin was performed with polyclonal anti-lamin (1:3000 dilution) and polyclonal anti-tubulin (1:3000), respectively, diluted in blocking buffer overnight at 4°C, and then incubated at room temperature for one hour with anti-rabbit-HRP conjugated antibodies at 1:2000 dilution with blocking buffer. The bands were visualized using the SuperSignal West Femto Maximum Sensitivity Substrate System (ThermoFisher Scientific). TLR2 immunoblotting was performed with anti-TLR2-specific polyclonal antibody (1:1000) and β-actin specific polyclonal antibody (1:3000).

### Mass spectrometry

IP was performed as described above. The bound immune complexes were eluted by heating in loading buffer, separated with 12% SDS-PAGE, and the gel stained with Coomassie. The gel pieces were de-stained in 200 μL of 25 mM ammonium bicarbonate in 50% v/v acetonitrile for 15 minutes and washed with 200 μL of 50% (v/v) acetonitrile. Disulfide bonds in proteins were reduced by incubation in 10 mM dithiothreitol (DTT) at 60°C for 30 minutes and cysteine residues were alkylated with 20 mM iodoacetamide (IAA) in the dark at room temperature for 45 minutes. Gel pieces were subsequently washed with 100 μL of distilled water followed by the addition of 100 mL of acetonitrile and dried on SpeedVac (Savant ThermoFisher). Then 100 ng of trypsin was added to each sample and allowed to rehydrate the gel plugs at 4°C for 45 minutes and then incubated at 37°C overnight. The tryptic mixtures were acidified with formic acid up to a final concentration of 1%. Peptides were extracted twice from the gel plugs using 1% formic acid in 50% acetonitrile. The collected extractions were pooled with the initial digestion supernatant and dried on SpeedVac (Savant ThermoFisher). Samples were desalted on Thermo Scientific Pierce C18 Tip.

Samples were analyzed on a LTQ Orbitrap Velos mass spectrometer (ThermoFisher Scientific) coupled to an Eksigent nanoLC-2D system through a nanoelectrospray LC-MS interface. An autosampler injected 8 μL of sample into a 10 μL loop. To desalt the sample, the material was flushed out of the loop, loaded onto a trapping column (ZORBAX 300SB-C18, dimensions 5×0.3 mm×5 μm), and washed with 0.1% formic acid at a flow rate of 5 μL/minute for 5 minutes. The analytical column was then switched on-line at 0.6 μL/minute over an inhouse made 100 μm i.d. × 200 mm fused silica capillary packed with 4 μm 80 Å Synergi Hydro C18 resin (Phenomex; Torrance, CA). After 10 minutes of sample loading, the flow rate was adjusted to 0.35 mL/minute, and each sample was run on a 90-minute linear gradient of 4–40% acetonitrile with 0.1% formic acid to separate the peptides. LC mobile phase solvents and sample dilutions used 0.1% formic acid in water (Buffer A) and 0.1% formic acid in acetonitrile (Buffer B) (Chromasolv LC–MS grade; Sigma-Aldrich, St. Louis, MO).

Data acquisition was performed using the instrument supplied Xcalibur™(version 2.1) software. The mass spectrometer was operated in the positive ionmode. Each survey scan of m/z 400–2000 was followed by collisionassisted dissociation (CAD) MS/MS of the twenty most intense precursor ions. Singly charged ions were excluded from CAD selection. Doubly charged and higher ions were included. Normalized collision energies were employed using helium as the collision gas.

MS/MS spectra were extracted from raw data files and converted into mascot generic format (mgf) files using a PAVA script (UCSF, MSF, San Francisco, CA). These .mgf files were then independently searched against the human SwissProt database (13188 entries) using an inhouse Mascot server (Version 2.6, Matrix Science). Mass tolerances were +/- 10 ppm for MS peaks, and +/- 0.6 Da for MS/MS fragment ions. Trypsin specificity was used allowing for 1 missed cleavage. Met oxidation, protein N-terminal acetylation, and peptide N-terminal pyroglutamic acid formation were allowed as variable modifications while carbamidomethyl of Cys was set as a fixed modification.

Scaffold (version 4.8, Proteome Software, Portland, OR, USA) was used to validate MS/MS based peptide and protein identifications. Peptide identifications were accepted if they could be established at greater than 95.0% probability as specified by the Peptide Prophet algorithm. Protein identifications were accepted if they could be established at greater than 99.0% probability and contained at least two identified unique peptides. All peptide sequences assigned are provided in **Supplementary Table S1**.

### Microscale thermophoresis

Microscale thermophoresis (MST) experiments were performed on a Monolith NT.115 Pico instrument (Nanotemper Technologies, München, Germany) to measure the binding affinity between AAT and the GR. AAT protein was fluorescently labeled using a Monolith Protein Labeling Kit RED-NHS 2nd Generation kit according to the manufacturer’s specifications. Recombinant human GR was serially diluted two-fold in nano-pure water two-fold from the stock concentration to create a 16-member concentration series (1.91 nM to 6.25 x 10^3^ nM). Fluorescently labeled AAT, prepared in PBS buffer with 0.05% P20, was added at a final concentration of 10 nM to the diluted GR fractions. Samples were then loaded into the Monolith NT.115 capillary tubes and loaded onto the Monolith NT.115 Pico instrument. For all the experiments, a Pico.RED detector was used at 20% laser power and medium MST power. Data were analyzed using the MO Affinity Analysis software suite (Nanotemper Technologies, Munich, Germany).

### Molecular modeling and docking

A model of GR was constructed using the known structures of DBD (PDB ID 5HN5) and LBD (PDB ID 1M2Z) domains and AlphaFold2 (Jumper *et al.*, 2021) to construct the hinge region connecting them. Structure refinement was performed using the *Rosetta Relax* protocol (Nivón *et al*, 2013). Given the intrinsic disorder of the NTD it was not modelled. Structural representations were produced using Schrodinger *PyMOL version 2.5.2*. Structural alignments were performed using *MUSTANG-MR* (Konagurthu *et al*, 2010). Docking simulations were performed using AlphaFold2 and ClusPro (Jumper *et al.*, 2021; Kozakov *et al.*, 2017).

### Isolation of nuclear and cytoplasmic fractions

Nuclear and cytoplasmic fractions of THP-1 macrophages were isolated using the NE-PER™ Nuclear and Cytoplasmic Extraction Reagents kit (ThermoFisher Scientific) according to the manufacturer’s instructions. Briefly, THP-1 macrophages were washed with 1X PBS and resuspended in cell extraction buffer supplemented with 1 mM PMSF and a protease inhibitor cocktail (Cell Signaling Technology (Danvers, MA). Nuclear and cytoplasmic protein fraction concentrations were quantified by the Bradford protein assay (Bio-Rad).

### Stable depletion of the glucocorticoid receptor (GR) in THP-1 cells

We employed shRNA-lentivirus technology to develop a pool of THP-1 cells stably depleted for GR. In brief, THP-1 cells were plated in 12-well tissue culture plates at a density of 1×10^4^ cells/well and infected with GR-directed shRNA or scrambled shRNA lentiviral particles at a multiplicity-of-infection of ≥1. THP-1 cells infected with the scrambled shRNA (THP-1 ^control^) and those stably depleted (knocked down, KD) for GR (THP-1^GR-KD^) were selected by adding 1 μg/mL puromycin dihydrochloride to the cell culture medium.

### RNA sequencing (RNAseq)

RNAseq and immunoblotting were used to confirm the depletion of GR mRNA and protein, respectively. For the former approach, we analyzed the expression of the *NR3C1* (GR) gene as part of another study of total RNA sequencing using the Illumina Next Generation Sequencing service from Genomics Shared Resource at the University of Colorado Cancer Center in THP-1^control^ and THP-1^GR-KD^ cells. RNAseq data were analyzed using DAVID Bioinformatics Resources 6.8 Vision software.

### p65 NFκB binding assay

The binding of the p65 subunit of NFκB was quantified using the TransAM™ NFκB p65 kit (Active Motif, Carlsbad, CA), as we previously reported (Bai *et al*, 2017).

### ELISA for interleukin-8 (IL-8) and angiopoietin-like 4 (ANGPTL4)

Control THP-1 and THP-1 GR knockout (as prepared above) macrophages were left untreated or treated with 1 μg/mL LPS alone or in the presence of 10 μM cortisol, 1 μM dexamethasone or 3 mg/mL AAT at 37°C in a 5% CO_2_ incubator. After 6 and 24 hours, the culture supernatants were quantified for IL-8 (R&D Systems) or ANGPTL4 (ThermoFisher Scientific) expression according to the manufacturer’s instructions.

### Infection of macrophages with *M. tuberculosis* and quantitation of cell-associated *M. tuberculosis*

*M. tuberculosis* infection of THP-1 macrophages and quantitation of cell-associated *M. tuberculosis* were performed as previously described (Bai *et al.*, 2017). In brief, differentiated THP-1 macrophages in 24-well tissue culture plates were infected with *M. tuberculosis* H37Rv at a multiplicity-of-infection (MOI) of 10 bacilli to 1 macrophage. For cells infected for one hour (Day 0), the supernatants were collected after one hour of infection, the cells washed twice with a 1:1 solution of RPMI:1X PBS, and the adherent cells lysed with 250 μL of a 0.25% SDS per well, followed by addition of 250 μL of 7H9 plating broth. Serial dilutions of cell lysates were prepared, and then 5 μL of each dilution was plated on Middlebrook 7H10 agar. For Day 2 and 4 infections, the cells were washed twice after the initial one hour of infection and replaced with RPMI medium containing 10% FBS. After two or four days of incubation, the supernatants were centrifuged to recover any non-adherent macrophages; these macrophages were lysed together with the adherent macrophages and *M. tuberculosis* cultured as described above.

## DATA AVAILABILITY

The mass spectrometry proteomics data have been deposited to the ProteomeXchange Consortium via the PRIDE [1] partner repository with the dataset identifier PXD030989 and 10.6019/PXD030989.

## Acknowledgment

We are grateful to Dr. Robert Sandhaus for helping to fund this research project. JRH is supported by NIH grants R03 AI139822 and R01 GM135421.

## Conflict of interest

The authors declare that they have no conflicts of interest with the contents of this article.

